# Association between COMT gene Val^158^Met heterozygote polymorphism and enhanced brain predicting processes

**DOI:** 10.1101/2020.09.26.314732

**Authors:** L. Bonetti, N.A. Sedghi, S.E.P. Bruzzone, N.T. Haumann, T. Paunio, K. Kantojärvi, M. Kliuchko, P. Vuust, E. Brattico

## Abstract

Predicting events in the ever-changing environment is a fundamental survival function intrinsic to the physiology of sensory systems, whose efficiency varies among the population. Even though it is established that a major source of such variations is genetic heritage, there are no studies tracking down auditory predicting processes to genetic mutations. Thus, we examined the neurophysiological responses to deviant stimuli recorded with magnetoencephalography (MEG) in 108 healthy participants carrying different variants of the Val158Met single-nucleotide polymorphism (SNP) within the catechol-O-methyltransferase (COMT) gene, which is responsible for the majority of catecholamines degradation in the prefrontal cortex. Our results showed significant amplitude enhancement of neural responses localized within inferior frontal gyrus, superior and middle temporal cortices to deviant auditory stimuli in heterozygote genotype carriers (Val/Met) vs homozygote (Val/Val and Met/Met) carriers. Integrating neurophysiology and genetics, this study provided new and broader insights into the brain mechanisms underlying optimal deviant detection.

## Introduction

Predicting the sensory environment is a fundamental animal function, depending on the tangled interplay between neurophysiology, genetics and biology. On a neurophysiological level, it is well-known that auditory predictions for sound environment are formed automatically in the supratemporal cortex and updated when errors are detected (Näätänen et al, 1978; Näätänen et al, 2007). These sensory processes have been tracked down with neurophysiological methods, giving rise to notorious components of the event related potential/field (ERF/P). Indeed, when a deviant stimulus is presented inserted in a sequence of coherent ones, the brain produces a negative response called mismatch negativity (MMN), which is usually followed by a positive component named P300. These events occur in a short time-window with a latency of about 100 to 350 ms from the onset of the deviant stimulus (Näätänen et al, 1978). Such components have been widely studied and provided several insights on how the brain detects and adapts to environmental irregularities (Näätänen et al, 2007). Indeed, MMN has been repeatedly connected to the predictive coding theory (PCT), which states that the brain is a constant generator of mental models of the environment that are progressively updated and refined on the basis of their match and mismatch with the external stimuli (Friston, 2012). Over the last decades, PCT has been successfully connected to the auditory domain and recently has been even adapted and explained in light of the peculiar and complex case of articulated music (Koelsch, Vuust, Friston, 2012). According to the PCT perspective, MMN has been considered an iconic evidence of the brain’s ability to make predictions of the upcoming events and automatically detect deviations from such predictions.

On an anatomical level, the originating brain sources of MMN have been especially located within the Heschl’s and superior temporal gyri, with a predominance of the right hemisphere (Garrido et al., 2009; Jemel et al., 2002; Rosburg et al., 2005). However, further studies also supported the existence of a functionally distinct and superordinate MMN generator in the frontal lobe (Deouell, 2007) which has been associated with the triggering of an involuntary attention switching process upon potentially critical unattended events in the acoustic environment (Giard et al., 1990; Näätänen et al., 2007; Rinne et al., 2000; Escera et al., 1998, 2003). Along this line, several studies have explored the connections between MMN and more complex cognitive abilities such as high-level attentive and memory processes and the correspondent individual differences among the population. For example, previous research has reported relationships between MMN amplitude and working memory (WM) (Bonetti et al., 2018) and sensory memory (Cheour, Leppänen, & Kraus, 2000). Furthermore, in a clinical review, Näätänen and colleagues (2012) has proposed MMN as a privileged brain response to index a variety of pathological states as well as individual differences related to healthy and impaired cognitive functioning.

Notably, even though it is established that a major source of such individual variations in cognitive processes is genetic heritage, there are no studies tracking down the brain’s auditory predicting processes to genetic mutations. Along this line, a great possibility comes from the recent advances in molecular and imaging genetics which opened up new opportunities to study auditory processing and cognitive functions. For instance, by combining the genetic variations of candidate selected genes with functional brain data broader insights may be achieved on the brain mechanisms regulating auditory predictive processing. In this light, a key candidate gene implicated in both physiological (Barnett et al., 2008; Berryhill et al., 2013; Bruder et al., 2005; Garcia-Garcia et al., 2017; Mattay et al., 2003) and pathological (de Diego-Balaguer et al., 2016; Hosák et al., 2007) neural conditions is COMT (Chen et al., 2004; Egan et al., 2001; Mier et al., 2010) (Bilder, Volavka, Lachman, & Grace, 2004), a gene coding for the catechol-O-methyltransferase enzyme which is responsible for the majority of catecholamines degradation in the prefrontal cortex (PFC) (Chen et al., 2004; Käenmäki et al., 2010; Matsumoto et al., 2003; Meyer-Lindenberg et al., 2005). Specifically, Val158Met (rs4680) is a common coding single-nucleotide polymorphism (SNP) involving the substitution of guanine (G) with adenosin (A) in the exon three of the gene that leads to a change in the amino acid located at position 158 of the codon. When G is not altered, a valine residual is coded in position 158 (G or Val allele). However, when it is mutated, valine is replaced by the evolutionarily more recent methionine (A or Met allele). Thus, three different COMT genotypes can be found within the population: Val/Val (A/A), Val/Met (A/G) and Met/Met (G/G) (Frank and Fossella, 2011; Männistö & Kaakkola, 1999, Mier et al., 2010; Meyer-Lindenberg et al., 2005). Notably, such change significantly affects enzyme activity and therefore the levels of prefrontal extracellular dopamine (Männistö, & Kaakkola, 1999). Specifically, the Val/Val version of the COMT enzyme breaks down dopamine with a degradation rate 40% faster than the Met/Met version (Chen et al., 2004; Lotta et al., 1995). Consequently, neurotransmitter is available at the synapses for extremely short periods in the case of Val/Val carriers, whereas it is preserved intact for a longer period in the case of Met/Met carriers (Barnett et al., 2008; Berryhill et al., 2013). The availability of neurotransmitter at the synapse largely influences neuronal activity, with both shortness and overplus of neurotransmitter undermining neuronal communication (Tritsch & Sabatini, 2012). In this regard, it has been suggested that an ideal dopamine availability seems to be maintained by the enzyme resulting from the heterozygous genotype Val/Met (Htun et al., 2014; Mazei et al., 2002; Schacht, 2016).

With regards to brain functioning, dopamine levels have been shown to be indissolubly related to brain activity (Barnett et al., 20018; Kähköen et al., 2002; Tritsch & Sabatini, 2012), and especially to PFC. Here, a large consensus has been established around the “inverted U-shaped” theory, stating that PFC requires an optimal, balanced level of dopamine and that higher or lower levels could result in impaired prefrontal mechanisms (Htun et al., 2014; Vijayraghavan et al., 2007; Stewart & Plenz, 2006). The interplay between COMT, dopamine and PFC has been highlighted by a positron emission tomography (PET) study which has supported the “U-shaped” model (Meyer-Lindenberg et al., 2005). In this light, the more frontal generators of the neural responses to deviant stimuli may represent an excellent opportunity to better understand the impact of COMT genetic variation on brain activity. Along this line, a previous electroencephalography (EEG) study with 22q11 Deletion Syndrome patients revealed an association between the homozygous COMT Met allele and a more marked speech MMN amplitude reduction in the PFC maximal amplitude channels (Baker et al. 2005). However, even though the study reported relevant results, it involved a clinical population of 50 participants and used only EEG measures.

Thus, in our study, using the combination of magnetoencephalography (MEG) and magnetic resonance imaging (MRI) in 108 participants, we investigated the relationship between COMT genetic variation and neural responses to deviant sounds in a healthy population. Specifically, we hypothesized to observe an enhancement of the MMN and P300 frontal generator amplitude in participant with COMT Val^158^Met heterozygote genotype (Val/Met) vs homozygote genotype (Val/Val and Met/Met).

## Results

### Sample Characteristics

The observed distribution of the different alleles in our sample was: Met/Met = 31(28.7%); Val/Met = 58(53.7%); Val/Val = 19(17.6%), coherently with the allele frequencies reported in previous studies (Frank & Fossella, 2011). According to ANOVAs and *X*^2^ tests, there were no significant differences among participants with respect to their COMT genotype, for age, sex, and handedness (**Table 1**).

**Table 1.** Demographic data of the participants illustrated according to their COMT genetic variation.

| Information | Whole sample | COMT Homo | COMT Etero | Homo vs Etero |
| --- | --- | --- | --- | --- |
| Participants | N 108 | N 50 | N 58 |  |
| Age | M 28.41, SE .771 | M 28.58, SE 1.16 | M 28.26, SE 1.05 | *$p > 0.05$ |
| Sex | M(N47, %43.5),<br>F(N61, %56.5) | M(N21, %42),<br>F(N29, %58) | M(N26, %44.8),<br>F(N32, %55.2) | **$p > 0.05$ |
| Handedness | A(1), L(6), R(101) | L(2), R(48) | A(1), L(4), R(53) | **$p > 0.05$ |
| *Independent samples Student *t*-test; **Chi-squared ($X^2$) test | | | | |

### COMT and neural responses to deviants

A 1000 permutation Monte Carlo simulation (MCS) was undertaken on the deviant neural responses, averaged for all the six sound deviants. As depicted in **Figure 1D**, the analysis revealed a single significant frontal cluster (*p* < 0.001, *k* = 260, time range from the onset of the deviants: 0.19 – 0.29 sec) where the neural amplitude was stronger for participants presenting the COMT heterozygote genotype (Val/Met) compared to homozygote genotype (Val/Val; Met/Met). Specific channels and time points are reported in **Table 2**. Additionally, the MCS conducted on single deviants identified several significant fronto-temporal clusters of channels. These differences were consistent across all deviants of the paradigm, peaking for pitch, slide and timbre, and are reported in **Table 3** and **Table ST1** and illustrated in **Figure 2**. Conversely, as expected the other direction of the contrast (COMT homozygotes vs heterozygotes) did not return any significant cluster for slide, timbre, rhythm, localization and intensity. In this case, we obtained a very small cluster for pitch only (*k* = 6; *p* < .001).

**Figure 1.**
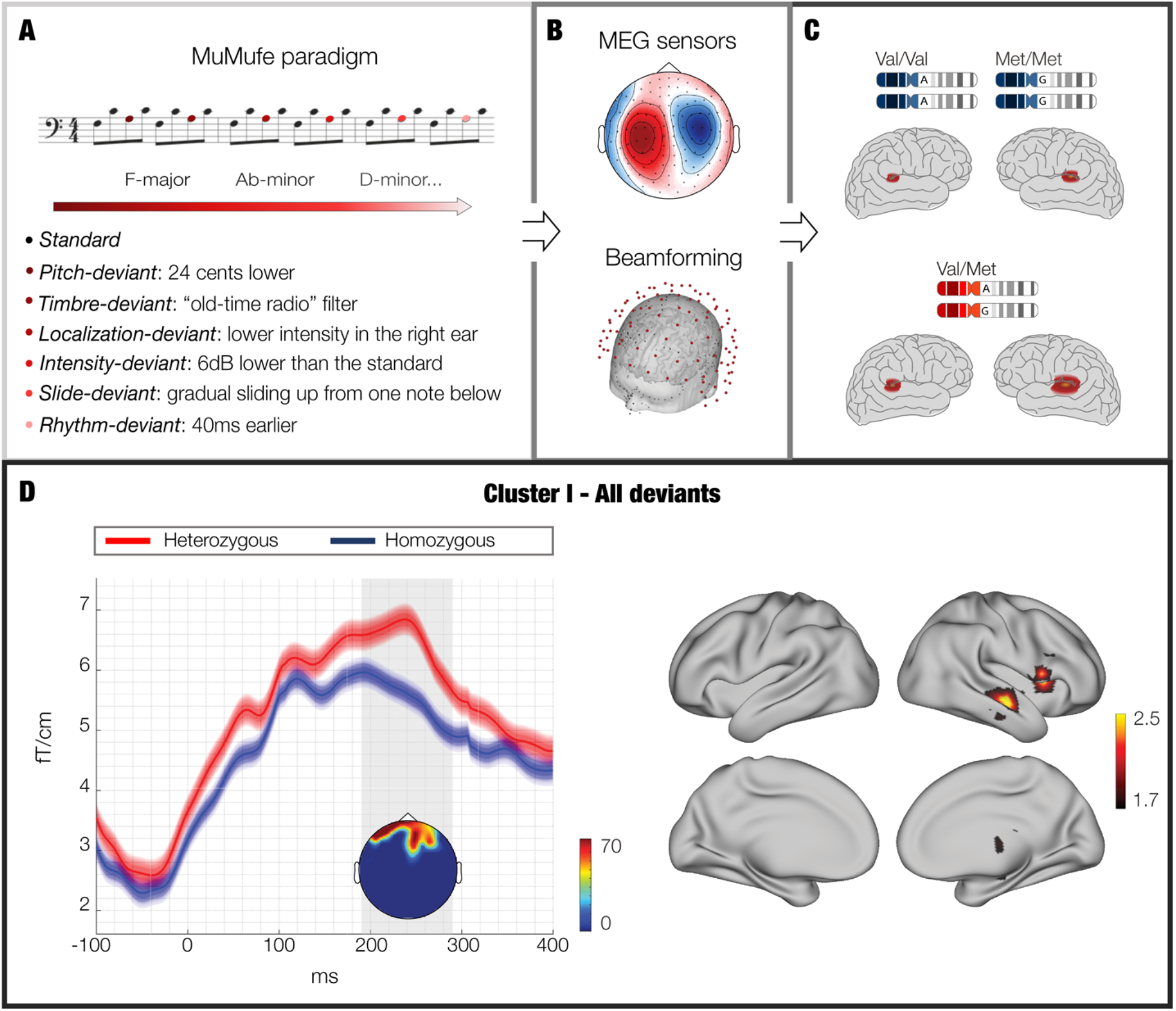
Overview of the analysis pipeline and main results. **A** – During MEG recordings, participants were presented with the musical multifeatures paradigm (MuMufe) while watching a silent movie. This paradigm allowed us to obtain the neural responses to deviant sound stimulations. **B** - MEG data has been collected, pre-processed and beamformed into source space. **C** – Participants have been divided according to their COMT genetic variation into two groups: heterozygotes (Val/Met) and homozygotes (Val/Val and Met/Met). **D** – Representation of the main significant cluster emerged by contrasting COMT heterozygote vs homozygote participants. Left figures show the MEG sensor results, while right ones provide a depiction of the brain sources originating the MEG signal.

**Figure 2.**
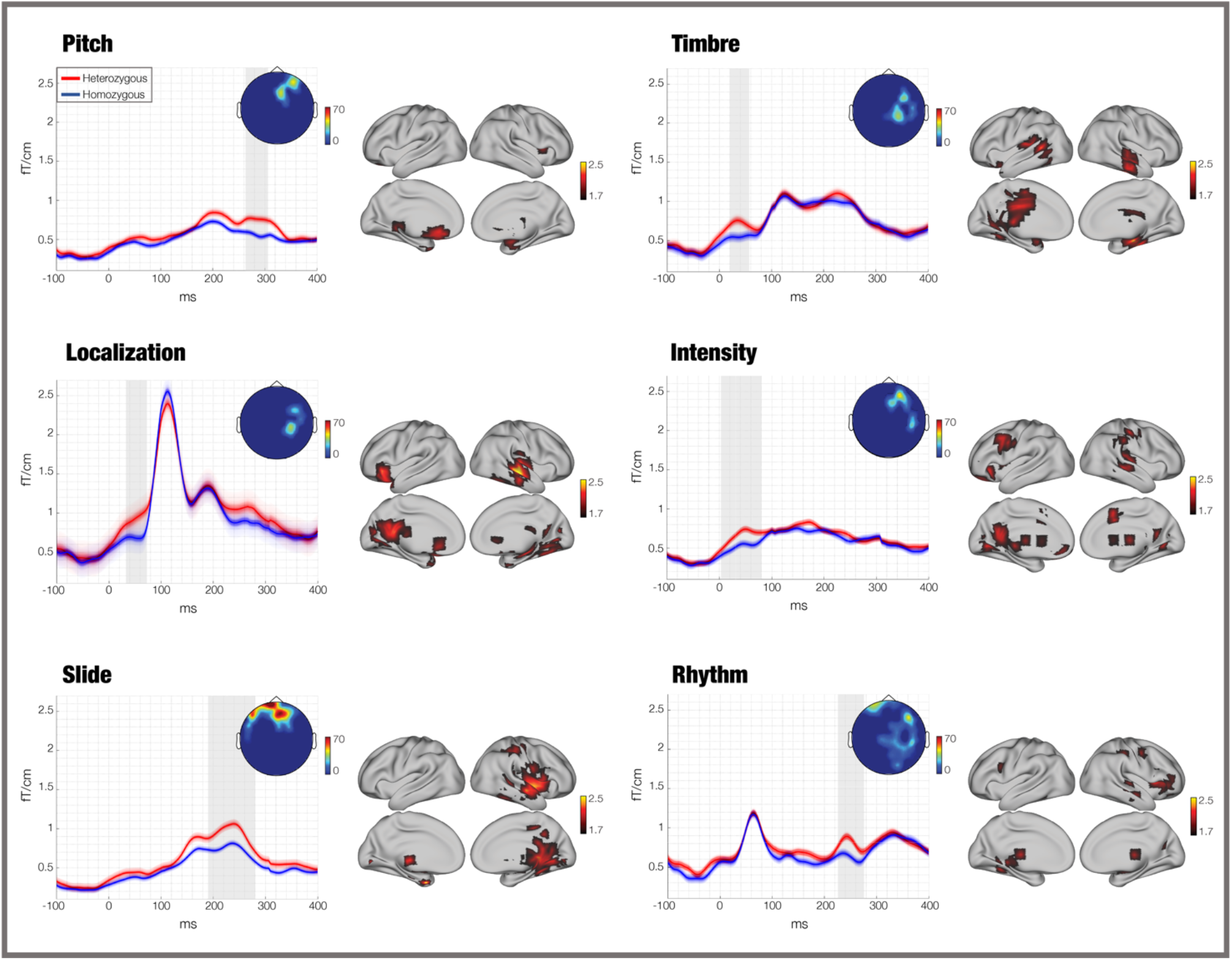
Neural responses to all deviants depicted according to the COMT polymorphism groups. Waveforms, topoplots and brain sources of the neural signal responses to all deviants depicted according to the COMT polymorphism groups in both MEG sensor and source spaces. Specifically, waveform images show the timeseries of the main significant cluster obtained contrasting the MEG sensor data of COMT heterozygotes vs homozygotes, while topoplots and source reconstruction plots report the spatial extent of those significant clusters, in MEG sensor and source space, respectively. With regards to topoplots, colorbars show the temporal extent (in ms) of the significant clusters, while source reconstruction plots colorbars illustrate the t-values of the contrasts.

**Table 2.** Significant channels and time-points of the cluster outputted by MCS on contrasts between the neural responses to all deviants of COMT heterozygote vs homozygote individuals.

| *MEG channels* | *Time (ms)* |
| --- | --- |
| 942 + 943 | 200 – 260 |
| 822 + 823 | 207 – 230 |
| 522 + 523 | 210 – 270 |
| 812 + 813 | 210 – 270 |
| 912 + 913 | 210 – 256 |
| 922 + 923 | 213 – 256 |
| 512 + 513 | 220 – 273 |
| 1022 + 1023 | 223 – 283 |
| 1222 + 1223 | 226 – 260 |
| 312 + 313 | 230 – 280 |
| 1212 + 1213 | 230 – 273 |
| 1232 + 1233 | 230 – 276 |
| 1012 + 1013 | 233 – 263 |
| 1032 + 1033 | 233 – 283 |
| 622 + 623 | 236 – 266 |
| 1042 + 1043 | 236 – 260 |
| 932 + 933 | 236 – 276 |
| 122 + 123 | 243 – 293 |
| 342 + 343 | 243 – 246 |
| 722 + 723 | 250 – 250 |
| 1112 + 1113 | 263 – 270 |
| 1242 + 1243 | 270 – 280 |

**Table 3.** Significant clusters outputted by MCS on contrasts between the neural responses to deviants of COMT heterozygote vs homozygote individuals. In this case, the analysis has been conducted independently for each of the six deviants. *k* refers to the spatio-temporal extent of the cluster (e.g. the overall number of channels and time-points forming the cluster).

| Deviant | Cluster # | k | MC simulations<br>*p* |
| --- | --- | --- | --- |
| Intensity | 1 | 85 | < .001 |
|  | 2 | 25 |  |
|  | 3 | 12 |  |
| Location | 1 | 53 | < .001 |
|  | 2 | 22 |  |
|  | 3 | 17 |  |
|  | 4 | 15 |  |
|  | 5 | 14 |  |
| Pitch | 1 | 84 | < .001 |
|  | 2 | 35 |  |
|  | 3 | 14 |  |
| Rhythm | 1 | 164 | < .001 |
| Slide | 1 | 263 | < .001 |
|  | 2 | 39 |  |
| Timbre | 1 | 82 | < .001 |
|  | 2 | 70 |  |
|  | 3 | 26 |  |
|  | 4 | 15 |  |
|  | 5 | 15 |  |

### Source localized activity

We reconstructed the brain sources of the MEG signal in the significant time-windows emerged from the previous MEG sensor analysis. To this aim, we used a beamforming approach and computed a general linear model (GLM) for each source reconstructed brain voxel and time-point. At the end, we corrected for multiple comparison with a cluster-based permutation test (Hunt et al., 2012). As depicted in **Figure 1D**, this analysis showed a stronger brain activity for Val^158^Met heterozygote vs homozygote participants mainly localised in the inferior frontal gyrus, and superior and middle temporal cortex. Source results for each of the six deviants are illustrated in **Figure 2**, while detailed statistical results for both averaged and single deviants concerning each brain voxel are reported in **Table ST2**.

## Discussion

In this study, we aimed to assess the relationship between different COMT genotypes and neural responses to deviant simulations. Our results showed a significant amplitude enhancement of such neural responses along the frontal MEG sensors in COMT heterozygote vs homozygote participants. Indeed, source reconstruction analysis located the neural sources concerning this difference especially within inferior frontal gyrus, and superior and middle temporal cortex.

The COMT polymorphism plays a unique role in modulating catecholamine flux and dopamine level in the PFC (Lewis et al., 2001; Mazei & al., 2002; Moron & al., 2002). Specifically, it has been suggested that an ideal dopamine level would be more frequently reached by the Val/Met carriers (Schacht, 2016), while Val/Val and Met/Met COMT variations would lead to a non-optimal dopamine degradation. As previous research showed, dopamine levels affect the extent of the brain activity, especially in the case of prefrontal regions. In particular, the “inverted U-shaped” theory states that PFC requires an optimal, balanced level of dopamine and that higher or lower levels can produce prefrontal impairment or deregulated activity (Egan et al., 2001; Htunet al., 2014; Joober et al., 2002; Vijayraghavan et al., 2007). Coherently with these evidences, our results showed that individuals with COMT heterozygote genotype (Val/Met) reported stronger neural amplitude than those with the homozygote genotype (Val/Val and Met/Met). As previously suggested by Schacht et al. (2016), this phenomenon may relate to the ideal dopamine degradation rate occurring in the COMT heterozygote individuals, that would be reflected in a stronger neural response to deviant sounds when compared to homozygote participants.

Additionally, our results are consistent with the findings reported by Baker et al. (2005). In a study with 22q11 Deletion Syndrome patients, they showed associations between the homozygous Met allele carriers and reduced speech MMN amplitude recorded with EEG. Our findings supported the results reported by Baker et al. (2005) and largely extended their significance. Indeed, here we showed a rather robust relationship between neural responses to deviants and COMT genetic variation in a large sample of 108 healthy participants, using a combination of MEG and MRI and a complex musical-multifeatures paradigm.

Furthermore, our beamforming analysis localized the sources of the difference between heterozygote and homozygote participants mainly within prefrontal areas such as inferior frontal cortex and secondary auditory regions as superior and middle temporal cortices. Notably, no differences were detected with regards to primary auditory cortex which is usually the main generator of the MMN.

This evidence suggests that SNPs in COMT gene may specifically affect the most frontal generators of the neural responses to deviants, regions whose activity is modulated by the dopamine level, that is largely regulated by COMT. Moreover, these frontal generators have been previously related to the involuntary attention-switching process towards critical unattended events suddenly occurring in the acoustic environment (Gallinat et al., 2003; Näätänen, Paavilainen, Rinne, & Alho, 2007). Thus, taken together, our results suggest a possible non-linear link between COMT Val^158^Met polymorphism, dopamine degradation and neural responses to deviants indexing automatic attentive brain processes. Moreover, our results can be extended to the superordinate framework represented by PCT, suggesting that to make adequate predictions and succesfully detect unexpected deviations the brain may require the optimal dopamine levels usually connected to the COMT heterozygote genotype (Val/Met).

In conclusion, our study showed the relationship between COMT Val^158^Met polymorphism and successfull deviant detection and automatic attentive functions, suggesting the relevance of investigating the complex interplay occurring between neurophysiology and genetics to better understand fundamental brain principles. Future research is called for to further explore the relationship between different COMT genotypes and brain activity and, to complement our work, especially focus on conscious attentional processes as well as the implications in pathological conditions.

## Materials and Methods

### Participants

This study was conducted within the large Tunteet protocol that involved the collection of neurophysiological, behavioral and genetical data of 140 participants. Previous results obtained from analysis of this dataset have been published in Bonetti et al. (2017), Bonetti et al. (2018), Alluri et al. (2017), Kliuchko et al. (2018); and Kliuchko, et al. (2016).

In this study, we considered the participants who took part in both the neurophysiological and genetical data gathering. The resulting sample consisted of 108 participants, 47 males (43.5%) and 61 females (56.5%), as reported in **Table ST1**. All participants were healthy, reporting no previous or current drug and alcohol abuse, were not under medication, did not report having had any neurological or psychiatric problems in their past, and declared to have normal hearing. Furthermore, their social, economic and educational status was homogeneous. The experimental procedures for this study were approved by the Coordinating Ethics Committee of the Hospital District of Helsinki and Uusimaa (approval number: 315/13/03/00/11, obtained on March the 11th, 2012). Moreover, all procedures were conducted in agreement with the ethical principles of Declaration of Helsinki.

### Stimuli and procedure

The stimuli adopted were piano tones from the Wizoo Acoustic Pianosample sounds from the software sampler Halion in Cubase (Stein- berg Media Technologies GmbH). The peak amplitude was normalized using Audition, Adobe Systems Incorporated. Because of its efficacy for balancing sounds on the basis on their most salient portion, peak amplitude normalization was used, labelled as the sharp attack. We employed the neurophysiological data elicited by the musical multi-feature paradigm (MuMUFE) because of its higher complexity with respect to other paradigms (e.g. oddball paradigm). Indeed, MuMUFE was widely showed to elicit clear MMNs and P3 in response to its deviants, and to be particularly effective for detecting also the prefrontal MMN and to study individual differences (Bonetti et al., 2018; Bonetti, Haumann, Vuust, Kliuchko, & Brattico, 2017; Vuust, Brattico, Seppänen, Näätänen, & Tervaniemi, 2012). The tones organization followed the common musical figure in Western music, known as “Alberti bass”, in patterns of four and with the arrangement in an arpeggiated chord (first–fifth–third–fifth). All the piano tones were of 200 ms in duration with 5 ms of raise and fall time. Interstimulus interval (ISI) was 5 ms. Every six patterns, the musical key of the presentation changed in pseudorandom order. The used keys were 24 (12 major and 12 minor) and were kept in the middle register. In each pattern, the third tone was replaced with a deviant of one of six types: pitch, timbre, location, intensity, slide and rhythm, as shown in **Figure 1A**. The deviant sounds were created by modifying one sound feature in Adobe Audition. The pitch deviant has been designed mistuning the third tone of the Alberti Bass by 24 cents, tuned downwards in the major mode and upwards in the minor one. To create timbre deviant, the ‘‘old-time radio” effect of Adobe Audition was applied to the sound. The location deviant was made by decreasing the intensity in one of the audio channels that resulted in perceptual shift of a sound source location from the center to a side. The intensity deviant was a reduction of a sound intensity by 6 dB. Slide deviant was made by gradual change of pitch from one note below up to the standard over the sound duration. The rhythm deviant was made by shortening a tone by 60 ms but keeping ISI of 5 ms, resulting in the consequent tone arriving earlier than expected. All the deviants were presented 144 times, half of which were played in a major and half in a minor mode, for an overall presentation of ~12 min. Randomization was conducted in Matlab and the stimuli were presented to the participants through Presentation software (Neurobehavioural Systems, Berkeley, CA). Participants were instructed to passively listen to sound sequences using headphones, Sennheiser HD 210.

Before the preparation for MEG recording, participants filled background questionnaire and performed a hearing threshold test utilizing the same sounds as in the experiment. We set the sound pressure level to 50 dB above the individual threshold. Then, participants were requested to watch a silenced documentary movie while comfortably sitting on a chair in a shielded chamber.

### MEG data acquisition

MEG data were collected at the Biomag Laboratory of the Helsinki University Central Hospital, in an electrically and magnetically shielded room (ETS-Lindgren Euroshield, Eura, Finland) with Vectorview™ 306-channel MEG scanner (Elekta Neuromag®, Elekta Oy, Helsinki, Finland). The scanner presented 102 sensor elements comprehending 102 orthogonal pairs of two planar gradiometer SQUID sensors and 102 axial magnetometer SQUID sensors. We placed the ground electrode on the right cheek, while the reference one was on the nose tip. Blinks, as well as horizontal and vertical eye movements, were measured with four electrodes attached above and below the left eye and close to the external eye corners on both sides. The sample rate of the registration was 600 Hz.

### MRI data acquisition

MRI data acquisition was conducted using a 3T MAGNETOM Skyra whole-body scanner (Siemens Healthcare, Erlangen, Germany) and a standard 32-channel head-neck coil. The measurements took place at the Advanced Magnetic Imaging (AMI) Centre (Aalto University, Espoo, Finland). We acquired 33 oblique slices every two seconds using a single-shot gradient echo planar imaging (EPI) sequence. The detailed parameters of the acquisition are reported as follows: field of view= 192 × 192 mm; 64 × 64 matrix; slice thickness = 4 mm, interslice skip = 0 mm; echo time = 32 ms; flip angle = 75°; voxel size: 2 × 2 × 2 mm3. We also collected T1-weighted structural images for individual coregistration. In this case, we used the following parameters: 176 slices; field of view = 256 × 256 mm; matrix = 256 × 256; slice thickness =1 mm; interslice skip = 0 mm; pulse sequence = MPRAGE.

### Pre-processing of MEG signals

First, aiming to minimize the affection of external and nearby noise sources and automatically individuate and correct bad MEG channels, the data was pre-processed by using Elekta Neuromag™ MaxFilter 2.2 (Elexta Oy, Helsinki, Finland) temporal Signal Space Separation (tSSS) (Taulu & Hari, 2009). We utilized the default inside expansion order of eight, outside expansion order of three, automatic optimization of both outside and inside bases, raw data buffer length of 10 seconds and subspace correlation limit of .98. The following data processing was performed by using FieldTrip, version r9093 (Donders Institute for Brain, Cognition and Behaviour/Max Planck Institute, Nijmegen, the Netherlands) (Oostenveld et al., 2011), and OSL (Woolrich et al., 2009), open source Matlab (MathWorks, Natick, Massachusetts) toolboxes widely used for MEG analysis. On average one channel (within the range 0 to 10 channels) per participant was marked ‘bad’ and replaced by interpolations of the activity measured in the neighbouring channels. The data was down-sampled from 600 to 300 Hz, and high- and low-pass filters were applied with cut-off frequencies at 1 and 25 Hz, respectively. Artefacts such as eye movements and cardiac activity were detected and removed by applying Independent Component Analysis (ICA) with the logistic infomax algorithm implemented in the *runica* function for Matlab (Haumann et al., 2016; Makeig et al., 1996). The number of removed artefactual ICA components per participant was on average .6 (range 1 to 3 components) for the MEG gradiometers and 2.6 (range 1 to 3 components) for the MEG magnetometers. Then, the data was segmented into epochs related to the six different deviant types and standard trials, choosing a pre-stimulus baseline correction from −100 to 0 ms in relation to the stimulus onset. We rejected trials with artefacts with amplitudes exceeding 2000fT, or 400fT/cm. The average of rejected trials was 2% for the MEG gradiometer data, evenly distributed across the deviant types and standard trials. Conversely, we did not discard any data for the MEG magnetometers. The average standard response of each participant was then subtracted from the correspondent average deviant responses to isolate the neural waveforms associated to deviant detection.

### Genotyping

Deoxyribonucleic acid (DNA) extraction was performed at the National Institute for Health and Welfare, Helsinki, Finland. DNA was extracted according to standard procedures. DNA samples were genotyped with Illumina Infinium PsychArray BeadChip and quality control (QC) was performed with PLINK. Markers were removed for missingness (>5%), Hardy-Weinberg equilibrium (p-value < 1 x 10-6), and low minor allele frequency (< 0.01). Individuals were checked for missing genotypes (>5%), relatedness (identical by descent calculation, PI_HAT>0.2) and population stratification (multidimensional scaling).

### COMT descriptive statistics

As illustrated in **Figure 1B**, participants were divided into two different groups, according to their COMT genotype. These groups were formed on the basis of the hypothesis that different COMT genotypes altered the optimal flux of catecholamines in the brain, and especially pre-frontal cortex (PFC) (on the “Inverted U” shaped model). Thus, COMT homozygotes Val/Val and Met/Met (altered level of catecholamines, lower and higher respectively) were grouped together into the homozygote group (50 participants), while Val/Met (optimal level) were grouped in the heterozygote group (58 participants). Differences between groups among demographic data and handedness were calculated through independent sample Student t-tests for continuous variables, and Chi-squared (*X*^2^) tests for categorical variables. Results are reported in **Table ST1**.

### MEG sensor brain responses to deviants and COMT

We performed statistical analysis only for MEG gradiometers because of their better signal-to-noise ratio compared to MEG magnetometers (in Bonetti et al. 2017, 2018, and Haumann et al., 2016 are presented quantitative measures of signal-to-noise ratio for this same dataset). As illustrated in **Figure 1C**, to test the hypothesized difference in terms of neural amplitude to deviants between the two COMT groups, a two-sample Student’s t-test was performed for each time point and each of the 102 gradiometer MEG channels. The differences were considered significant with *p* < .01. The t-tests were conducted in a 300-ms time-window from the sounds onset with a sampling rate of 300 Hz, resulting in 92 time-points (~ 3.33 ms each). To correct for multiple comparisons, a 1000-permutation MCS was computed to identify the significant clusters of neighbouring channels where the neural amplitude differed between the two COMT groups. First, we conducted this procedure on the neural responses averaged for the six deviants. Afterward, in order to deepen the analyses, we performed the same procedure independently for the neural response to each deviant. We considered significant the clusters that emerged from the MCS with a cluster-forming threshold of *p* < .0014 (corresponding to α = .01 divided by the seven independent analyses that we performed). Finally, we calculated analogous MCS also on the other direction of the contrast (COMT homozygotes vs heterozygotes).

### Source reconstruction and COMT

As depicted in Figure 1B, after computing the statistical analysis for MEG sensors, we conducted a further investigation in source space. To this aim, using OSL, a freely available toolbox for MEG analysis (Woolrich et al., 2009), we reconstructed the sources of the MEG signal recorded on the scalp by using an overlapping-spheres forward model and a beamformer algorithm as inverse model (Hillebrand & Barnes, 2005). We used an 8-mm grid including both planar gradiometers and magnetometers. Specifically, the spheres model was an approximation of the MNI-co-registered anatomy, represented as a simplified geometric model that used a basic set of spherical harmonic volumes (Huang, Mosher, & Leahy, 1999). Conversely, the beamforming algorithm utilized a set of weights that were sequentially applied to the source locations to estimate the contribution of each brain source to the activity recorded by the MEG sensors. This was done for each time-point (Brookes et al., 2007; Hillebrand & Barnes, 2005). Then, we contrasted the brain activity reconstructed in source space in response to deviant vs standard sound stimulation. This was done by using a GLM at each dipole location and for each time-point (Hunt et al., 2012). Afterwards, we calculated the absolute value of the reconstructed time-series to prevent sign ambiguity of the neural signal and we computed first-level analysis, consisting of contrasts of parameter estimates for each time-point, dipole and participant. These results were then submitted to a second-level analysis, using paired-sample t-tests contrasting COMT heterozygote vs homozygote participants. Here, we employed a spatially smoothed variance computed with a Gaussian kernel (full-width at half-maximum: 50 mm). In conclusion, to correct for multiple comparisons, we utilized a cluster-based permutation test (Hunt et al., 2012) with 5000 permutations on the second-level analysis results. In this case, we investigated only the significant time-windows emerged from MEG sensor level analysis independently for each deviant and therefore we considered an a level = .05, corresponding to a cluster forming threshold *t*-value = 1.7.

## Acknowledgements

We thank the entire team who contributed to the “Tunteet” neuroscientific and psychological data collection: Brigitte Bogert, Benjamin Gold, Johanna Normström, Taru Numminen-Kontti, Mikko Heimölä, Toni Auranen, Marita Kattelus, Jyrki Mäkelä, Mikko Sams, Petri Toivainen, Mari Tervaniemi, Anja Thiede, Alessio Falco, David Ellison, Chao Liu, Suvi Lehto and Simo Monto.

The Center for Music in the Brain (MIB) is funded by the Danish National Research Foundation (project number DNRF117).

We would also like to acknowledge the financial support of the Academy of Finland (project number 133673), the 3-years grant of the University of Helsinki and the three-month research stay grant to assess potential PhD candidates offered by Aarhus University Foundation to Nader Alessandro Sedghi.

## Competing interests statement

The authors declare no competing interests.

## Data availability

The code and multimodal neuroimaging data from the experiment are available upon reasonable request.

## Author contributions

EB, LB, MK, TP, KK conceived the hypotheses and designed the study. LB, NTH, NAS performed pre-processing and statistical analysis. EB, PV, SEPB, TP and KK provided essential help to interpret and frame the results within the neuroscientific literature. NAS and LB wrote the first draft of the manuscript. SEPB and LB prepared the figures. All the authors contributed to and approved the final version of the manuscript.

## SUPPLEMENTARY MATERIALS

As follows, supplementary tables related to this study. In the cases when the supplementary tables were too large to be conveniently reported in the current document, they have been reported in Excel files that can be found at the following link: https://www.dropbox.com/s/jv8tuq4kpi5d39f/TableST2_COMTDeviants.xlsx?dl=0

**Table ST1.**
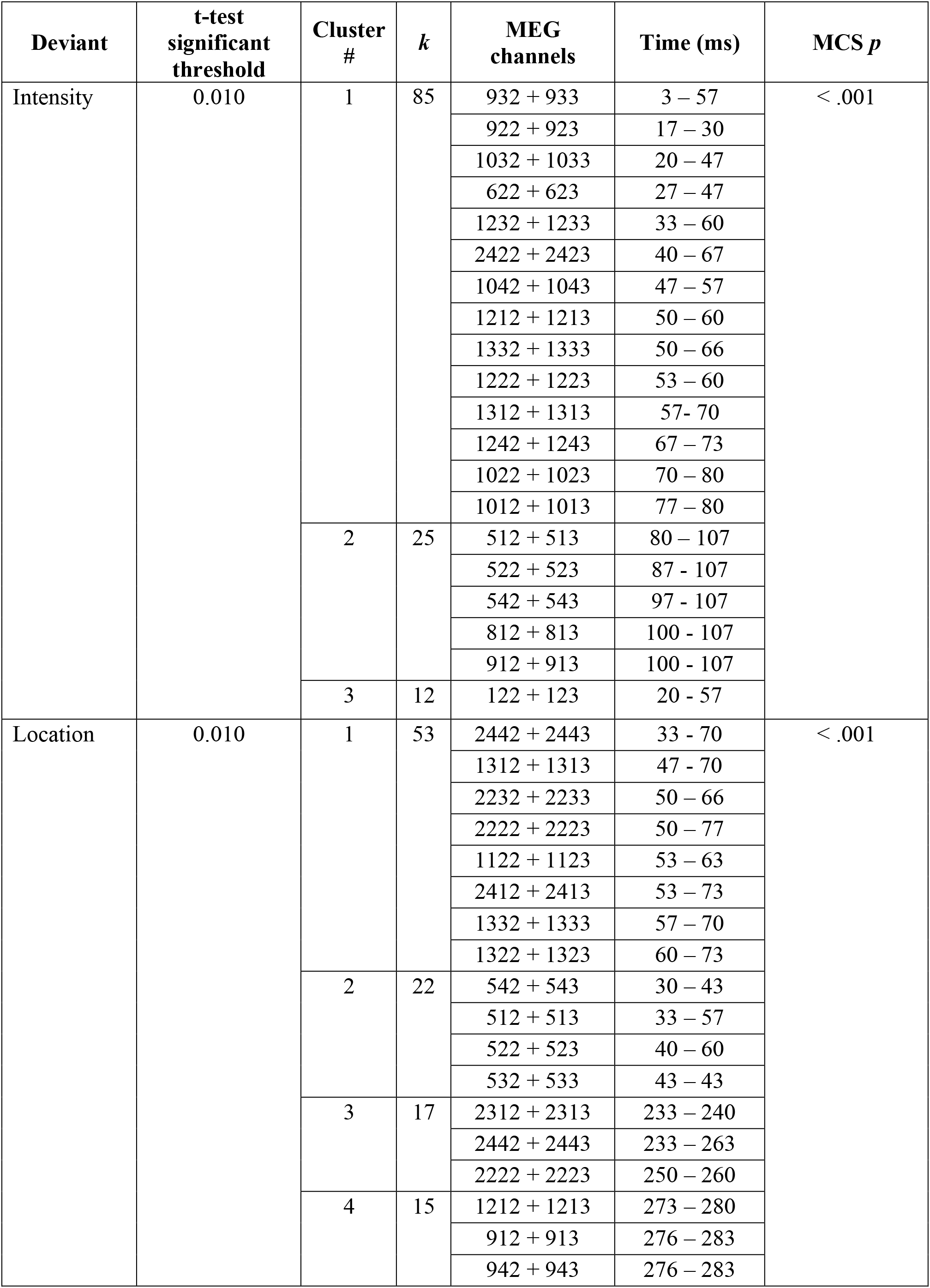
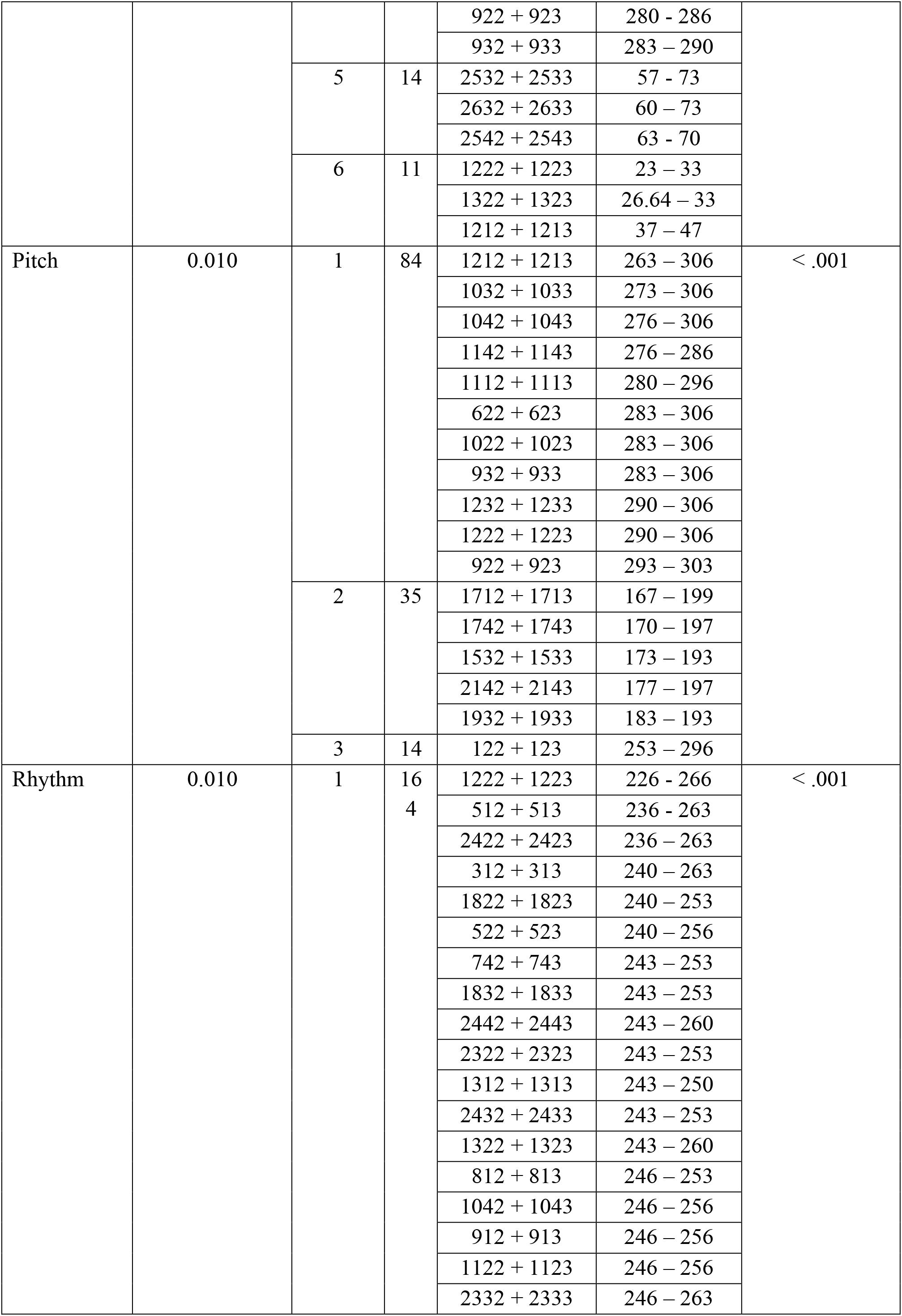
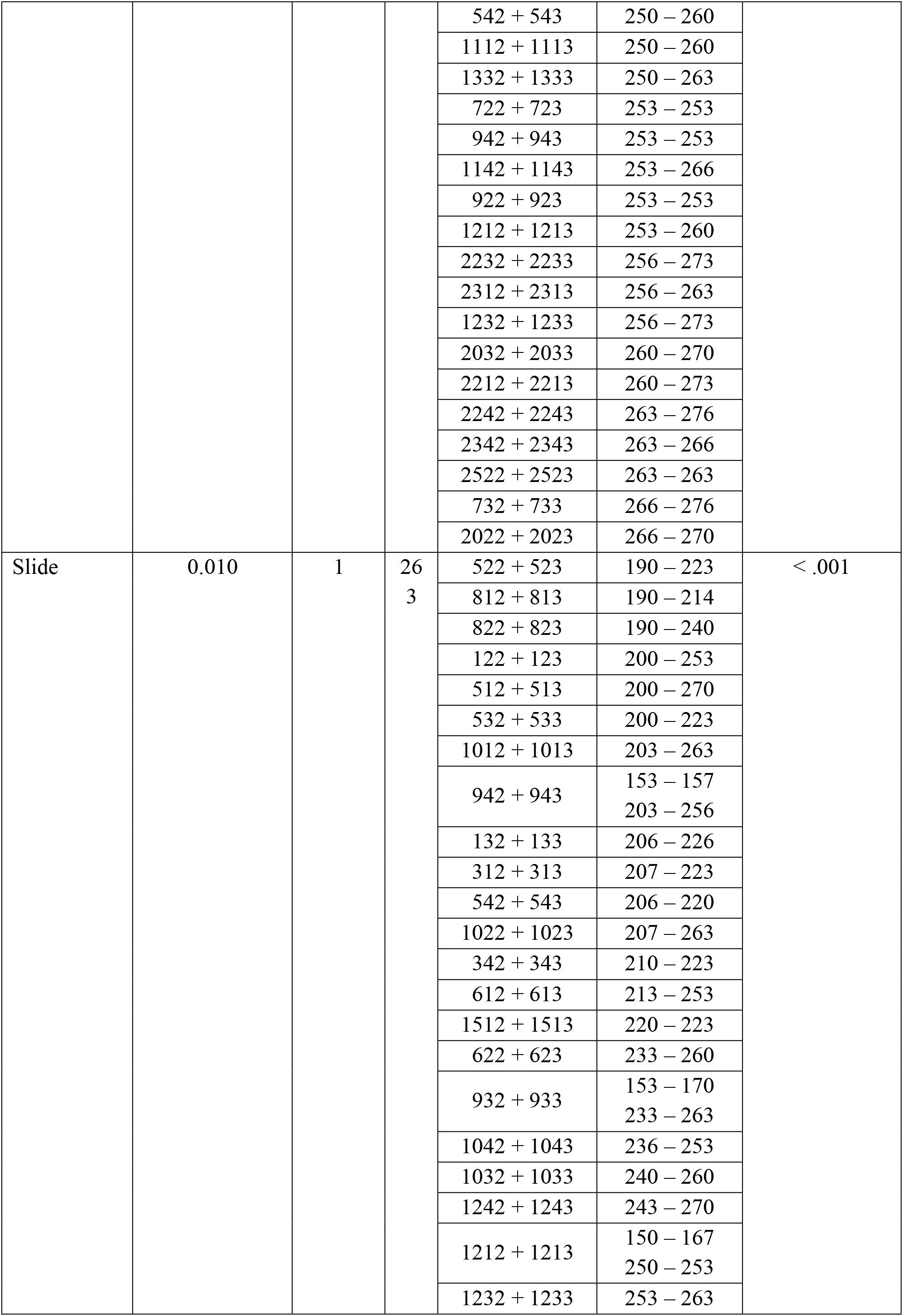
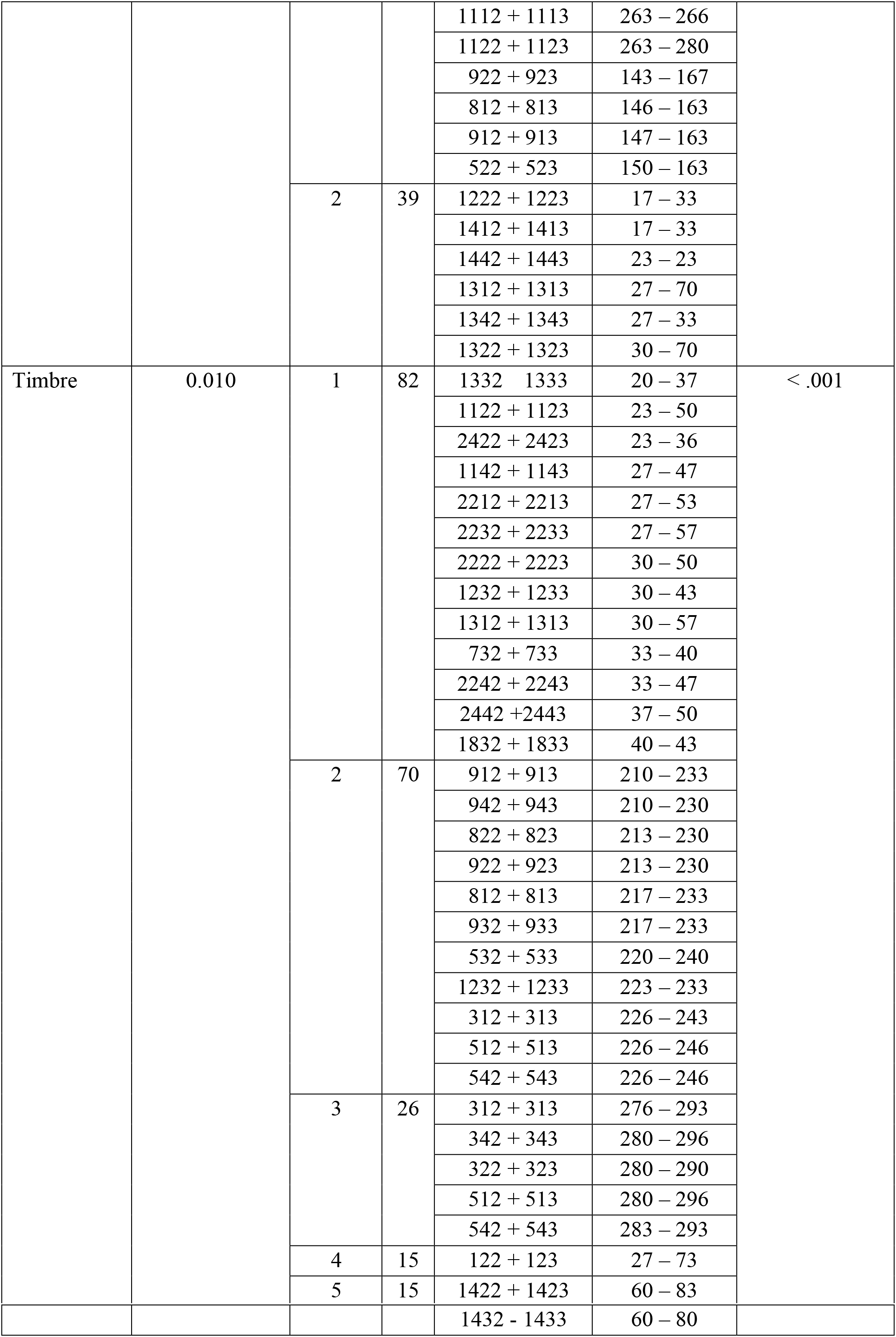
Detailed information on the significant clusters outputted by MCS on contrasts between the neural responses to deviants of COMT heterozygote vs homozygote individuals. In this case, the analysis has been conducted independently for each of the six deviants. *k* refers to the spatio-temporal extent of the cluster (e.g. the overall number of channels and time-points forming the cluster).

***Table ST2 – Detailed information of significant MEG source clusters for the contrast between COMT heterozygote vs homozygote participants***

Significant clusters of MEG sources emerged from MCS contrasting COMT heterozygote vs homozygote participants. The excel file depicts those clusters with regards to significant voxels, timewindows and averaged *t*-values for each voxel. This information is reported for the average of all deviants and for the six deviants independently.

## Notes

### Competing Interest Statement

The authors have declared no competing interest.

